# Predicting Response to McGurk Illusion Based on Periodic and Aperiodic Prestimulus EEG Activity

**DOI:** 10.1101/2022.01.20.477172

**Authors:** Vinsea A V Singh, Vinodh G Kumar, Arpan Banerjee, Dipanjan Roy

## Abstract

Studies have reported that prestimulus brain oscillations guide perceptual experiences during AV speech perception. However, ’what’ features of these prestimulus activity drives perception remains unknown. In this electroencephalography (EEG) study, we investigated the relationship between prestimulus periodic oscillations and aperiodic components with subsequent perception of the McGurk illusion at single-trial levels. Using logistic mixed-effect models, we determined which spectral features from different sensor regions predicted response to the illusory perception. We found that lower alpha (8-12 Hz) and beta (15-30 Hz) band oscillations over parieto-occipital sensors predicted illusion perception. We also found lower aperiodic offset values over parieto-temporal sensors, and lower ‘global’ effect of exponent over the scalp, that predicted the response to McGurk illusion. We conclude that the predominant source of the prestimulus oscillatory state is aperiodic background activity, and that variations in these arrhythmic components account for inter-trial and inter-individual variability in perception of the McGurk illusion.

## 1. Introduction

Perception is driven by both incoming stimuli and the functional state of the brain (Jensen *et al*., 2012). The extent to which the functional state of the brain impact conscious perception has been an area of intense investigation, and growing evidence indicates that ongoing/spontaneous neural activity constitutes an essential property of the neural network that underlies cognition and is related to variability in perception (Dijk *et al*., 2008; Kayser *et al*., 2016; Spadone *et al*., 2020). Nevertheless, the functional relevance of the ongoing neural activity before speech perception remains poorly understood.

Speech perception, especially during face-to-face conversation, involves temporal integration of the auditory and visual inputs, requiring processing across a broadly distributed cortical network. Consequently, fluctuations in neural activity within these networks become a potential source of variability in perception. While extensive prior work has characterized the contextual and stimulus-related factors that contribute to the variability in speech perception (Olasagasti & Giraud, 2020; for review, see Gagnepain *et al*., 2021), the impact of multisensory speech perception as a function of the subjective difference in the ongoing neural activity (or prestimulus activity), remains unclear. Studies on multisensory speech perception have predominantly employed the McGurk effect, wherein participants report an illusory/fused percept when presented with a mismatching AV stimulus (for example, an auditory /*ba*/ dubbed over the visual lip movement of /*ga*/ giving rise to an illusory percept of /*da*/, McGurk & MacDonald, 1976). These studies have also shown that the probability of the occurrence of the McGurk effect is not equal across participants, and also illusion is not experienced in every trial (Nath & Beauchamp, 2011, 2012; Mallick *et al.,* 2015). This makes the McGurk paradigm a lucrative model to investigate the impact of ongoing neural activity in effectuating subsequent perception. Keil *et al.,* 2012 have previously reported that an increased beta-band power in the left superior temporal gyrus (lSTG) prior to the McGurk trial result in a fused percept. While the aforementioned evidence substantiates the functional relevance of ongoing neural oscillations in orchestrating illusory perception, the study does not account for the inter-individual differences in the susceptibility of the McGurk effect. Also, their study has estimated these power fluctuations in the prestimulus duration for the averaged trials, thereby overlooking neuronal fluctuations on a trial-by-trial basis, which would give a better insight regarding perceptual variability across trials.

Moreover, studies suggest that the neural oscillations one computes and compares with the cognitive, perceptual, and behavioural states do not directly reflect the rhythmic activity (Bullock *et al.,* 2003; Buzsáki *et al*., 2013; He B., 2014; Donoghue *et al*., 2020; Wen & Liu, 2016). Rather, in the frequency domain, oscillations manifest as narrowband peaks of power over and above the non-oscillatory (or aperiodic) component. This arrhythmic brain activity has been linked to be modulated in different cognitive, perceptual, and behavioral states (Engel *et al*., 2001; Buzsáki & Draguhn, 2004; He B. *et al*., 2010; Podvalny *et al*., 2015). So, the changes in power observed might be because of changes in the aperiodic component, even when no periodic oscillation is present. Due to this, systematic parameterization of the power spectrum is required to circumvent misinterpretation of the observed narrowband power differences between conditions of interest (Cole and Voytek, 2019; Lansbergen *et al*., 2011).

Overall, in this study we aimed to first investigate the changes in periodic and aperiodic oscillations before the incongruent McGurk trials. We looked at differences in prestimulus parameterized power when the individuals perceived versus not perceived the McGurk illusion, thereby, capturing between subject variability. We then examined the differences in the prestimulus periodic and aperiodic parameters between illusory and non-illusory trials for different sensor regions. This was done to understand the distribution of power across the whole brain. And finally, to understand which spectral features significantly predicted the behavioral response trail-wise, independent of inter-subject variability, we employed a logistic mixed effect model where the dependent variable was the binary perceptual response (illusion or no-illusion), and the periodic and aperiodic parameters from different sensor regions were the predictors. The model also recognized the relationship between brain oscillations prior to the stimulus onset and subsequent McGurk percept on a trial-by-trial basis. For the current study we reanalyzed EEG data collected in a previous study that explained using a biophysical model, the underlying mechanisms governing large-scale brain network dynamics between rare and frequent groups of McGurk perceivers (Kumar *et al*., 2020). To the best of our knowledge, this is one of the first studies to pinpoint how specific prestimulus neural differences in both aperiodic and oscillatory components index the veracity and experience of multisensory illusory speech perception.

## 2. Materials and Methods

### 2.1. Participants

For this study, we have used the previously recorded EEG and behavioural data described in Kumar *et al*. (2020), where eighteen healthy right-handed participants (8 females) with a mean age of 24.9 ± 2.8 years were recruited for the study. A written informed consent was obtained from the participants under the experimental protocol approved by the Institutional Human Ethics Committee (IHEC) of the National Brain Research Centre (NBRC), India, in accordance with the 2008 Declaration of Helsinki.

### 2.2. Stimulus Design and Trials

Participants completed a two alternate forced-choice task, where they reported their subjective perception to the audio-visual (AV) stimuli as described in Kumar *et al*. (2020). Each participant was subjected to four kinds of AV stimuli: three congruent (audio syllable matching with the video articulation) syllables */pa/*, */ta/*, and */ka/*; and one McGurk (audio-visual mismatch) syllable (auditory */pa/* with visual */ka/*) producing the illusion of syllable */ta/*. As the participants observed the four stimuli presented at random, they reported if they heard either /*pa*/, /*ta*/, /*ka*/, or "something else," being unaware of the McGurk illusion. The experiment was carried out for five blocks, where each block consisted of 120 trials (30 trials of each AV-stimuli presented at random). Inter-stimulus intervals were pseudo-randomly varied between 1200ms (milliseconds) to 2800ms to minimize expectancy effects (Minor, 1970).

### 2.3. Data acquisition and pre-processing

A Neuroscan EEG recording and acquisition system (SynAmps2, Compumedics, Inc.) with 64 Ag/AgCl scalp electrodes molded on an elastic cap in a 10-20 montage was used to collect continuous EEG scans, where Cz was made the reference electrode. The sampling rate was 1000 Hz, and the channel impedances were kept below 10kΩ. The FASTRAK 3D digitizing system was used to register individual electrode locations (Polhemus Inc.).

The continuous EEG data collected were preprocessed using EEGLAB toolbox (Delorme and Makeig, 2004) and custom MATLAB codes (version R2020a). A finite impulse response (FIR) filter was applied at 0.1 Hz and 80 Hz to the data, followed by a 9th order 2-pass Butterworth filter (notch-filter) between 46 and 54 Hz to remove the line noise. The data was then average re-referenced. Eye blinks and muscle artifact components from the signal were removed by independent component analysis (ICA) using the *runica* function. Epochs of 0.8s (-0.8s to 0s) before the stimulus onset (prestimulus) and 0.8s post the onset of the stimulus (post-stimulus) were extracted using the trigger information (**Figure 1A**). The extracted epoch data were sorted based on the Congruent AV stimuli (/pa/, /ta/, and/ka/) and the incongruent McGurk stimulus. The sorted prestimulus and post-stimulus epochs were then baseline corrected by removing the temporal mean of the EEG signal on an epoch-by-epoch basis. Finally, to remove response contamination from any other artefacts, epochs with amplitudes above and/ or below ±75 µV were removed from all electrodes.

**Figure 1:**
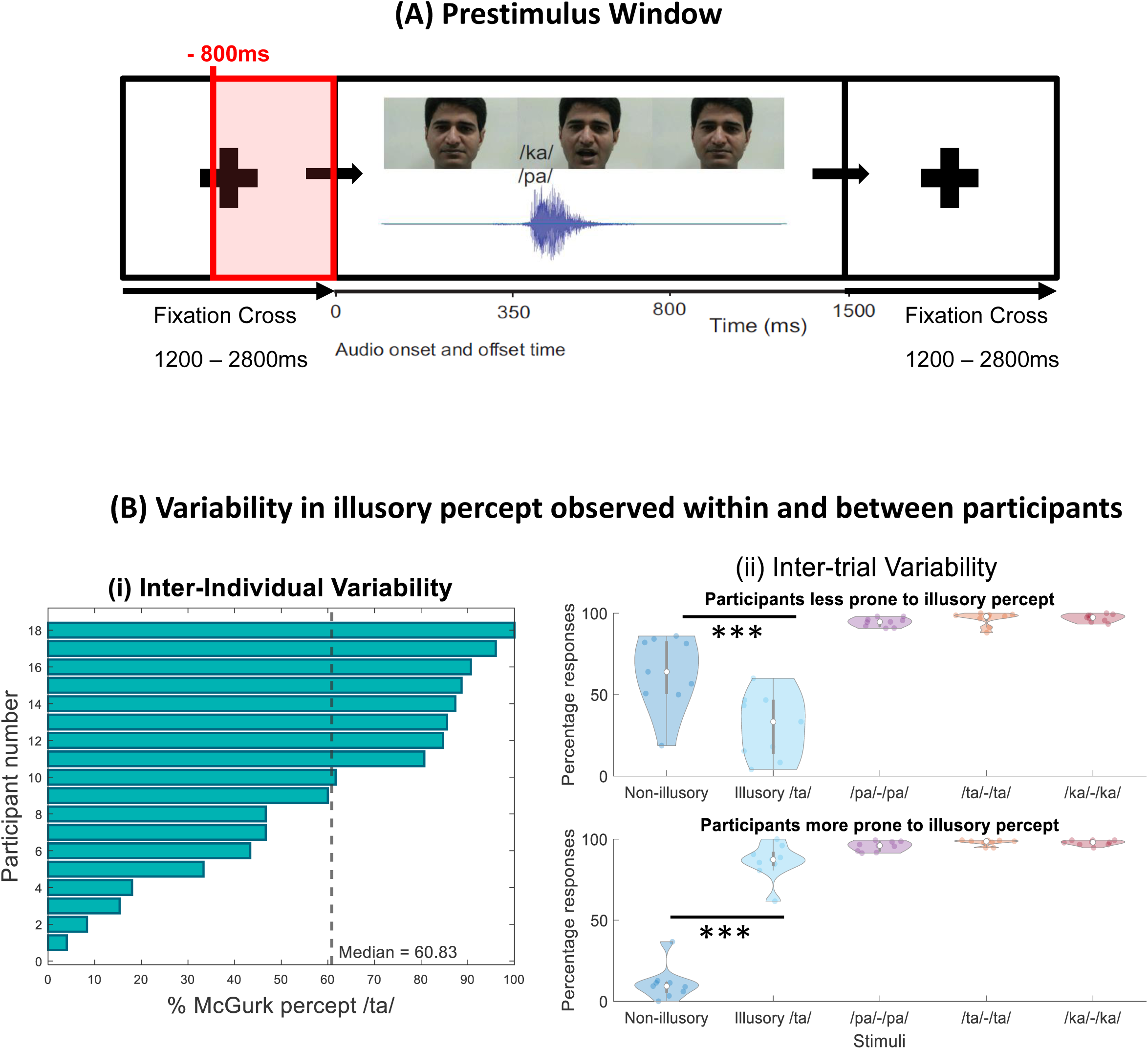
(A) Prestimulus time-window. Example trial with a fixation cross pseudo-randomly varied between 1200 – 2800ms before the onset of the AV stimulus. The red box indicates the time window chosen for prestimulus epoch data of 800ms duration. **(B) Behavior.** (i) Bar graph representing inter-individual variability - Propensity of McGurk effect for all the 18 participants expressed as the percentage of */ta/* percept during the presentation of the McGurk illusion. (ii) Violin plots showing inter-trial variability - Percentage of */ta/* (illusory) and */pa/* (non-illusory) percept during the presentation of McGurk stimulus and the congruent AV stimuli (*/pa/, /ta/,* and */ka/*) averaged over participants less prone (< median percentage response) and more prone (> median percentage response) to the illusion. The white dot in the center of each violin plot represents the median.

### 2.4. Spectral analysis

We computed the spectral power at each electrode on a trial-by-trial basis using the multi-taper Fast Fourier Transform (mtfft) method, for both the prestimulus and the post-stimulus epoch data. Power spectra were computed to extract the distribution of signal power over different frequency bands (theta (4-7 Hz), alpha (8-12 Hz), beta (15-30 Hz), and gamma (31-45 Hz)) for all the stimulus conditions (incongruent McGurk, and congruent) using the Chronux toolbox function *mtspectrumc.m* (Bokil *et al*., 2010) and customized MATLAB codes. Time bandwidth product and the number of tapers used were set to 3 and 5, respectively.

### 2.5. Extracting the periodic and aperiodic components from the power spectral densities (PSDs)

To separate the "background" *1/f* aperiodic component from its periodic counterpart, we used FOOOF (or Fitting Oscillations and One Over *f*) algorithm (Donoghue *et al*., 2020). FOOOF algorithm takes the original power spectral densities (PSDs) and extracts the aperiodic signal, and superimposes them on periodic oscillatory components, referred to as "peaks." These peaks are considered to be oscillations and are modelled individually as Gaussian functions. Each of these Gaussian has three parameters that are used to define an oscillation: center frequency (CF) which is the mean of the Gaussian, amplitude of the peak which is the distance between the peak of the Gaussian and the aperiodic fit (PW), and bandwidth (BW) as two standard deviations. The aperiodic component is defined by two parameters: exponent or negative slope (Miller *et al.,* 2009), and offset which is y-intercept across frequencies (Donoghue *et al.,* 2020).

The periodic and the aperiodic components were extracted from the prestimulus and poststimulus power spectra of each electrode trial-wise using the MATLAB implementation of FOOOF (version 1.0.0) which is originally written in python by Donoghue *et al*., 2020. Power spectra were parameterized across the frequency range 0.1 to 45 Hz using customized MATLAB scripts. The periodic oscillations were extracted by subtracting the aperiodic fit from the original PSDs.

### 2.6. Predicting response to the McGurk trials from prestimulus parameterized power

We were interested to assess the relationship of prestimulus periodic and aperiodic parameters from each sensor over participants’ overall responses to the upcoming McGurk trials, and predict the response based on prestimulus spectral features. To estimate that, a logistic mixed model was fitted using the *glmer* function in R (R Development Core Team, 2023). Where, two possible behavioral outcomes (i.e., illusion vs. no-illusion) were chosen as the dependent variable. The periodic (center frequency, aperiodic adjusted power, and bandwidth) and aperiodic (offset, exponent) parameter predictors, computed across different sensor regions (frontal, central, parietal, occipital, and temporal), were chosen as fixed effects. Since, we were interested to determine which features could predict the behaviour, our model assumed that the effect of each predictor on dependent variable is independent of other predictors in the model. Moreover, to account for the between-subject variability, we used subject ID as the random effect. The fixed effects were mean-centered around zero (Hox, 2002), and the random effect model selected was based on the lowest Akaike information criterion (AIC) and Bayesian information criterion (BIC) value based on the model’s log-likelihood ratio (Field *et al*., 2013). The formula was defined as follows:

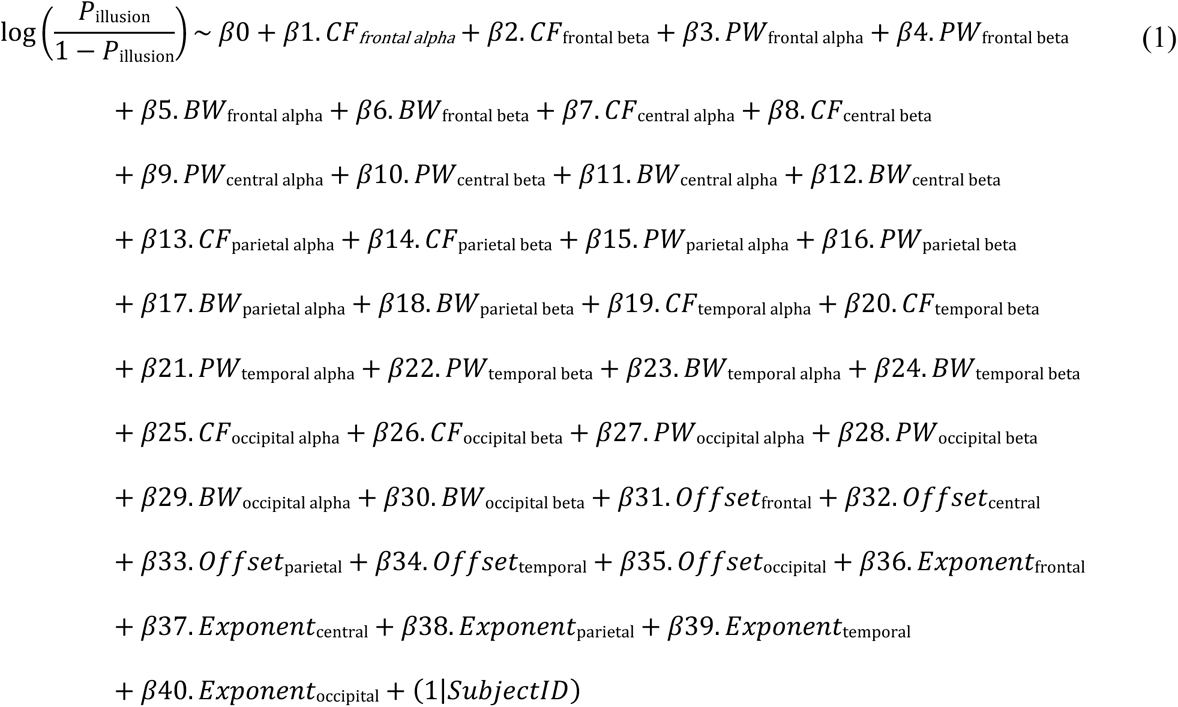

Where, P_illusion_ is the probability of perceiving the illusion, β_0_ is the intercept, β_n_ are regression weights. The predictors included are the CF which is the center frequency, PW is the peak parameter, and BW is the bandwidth across alpha (8-12 Hz) and beta (15-30 Hz) frequency bands, and across different brain regions (Regions: frontal, central, parietal, occipital, and temporal). These continuous variables define the periodic component of the power spectra. The aperiodic component was defined similarly by offset and exponent. Standardized parameters were obtained by fitting the model on a standardized version of the dataset, where the significance of each beta coefficient was tested against zero (i.e., Bn = 0). The 95% Confidence Interval (CI), and p-values were computed using a Wald z-distribution approximation (Demidenko, 2020).

Furthermore, to quantify the evidence of each predictor in the estimated model, we computed the Bayes factor (BF) of each independent predictor using the *BayesFactor* library in R (Morey and Rouder, 2015). The Bayes factor was calculated by comparing the full model to models in which the given predictor was removed, indicating the effect of the given predictor on the model fit in predicting the response. A Bayes factor value of less than one (BF<1) indicated that the removed predictor improved the overall model fit, whereas a Bayes factor value greater than one (BF>1) showed little to no improvement in model fit without the given predictor.

Finally, after estimating significant factors that predicted the trial-wise response to the upcoming McGurk stimulus, we also fitted a logistic mixed effect interaction model for the periodic and aperiodic parameters, with Subject ID as the random effect. The model estimated the effects of one predictor on response depending on the value of another predictor and vice-versa (Fisher, 1992). The interactions were estimated between periodic parameters of alpha and/ or beta frequency bands, and also between aperiodic parameters. The formula applied for the model was as follows:

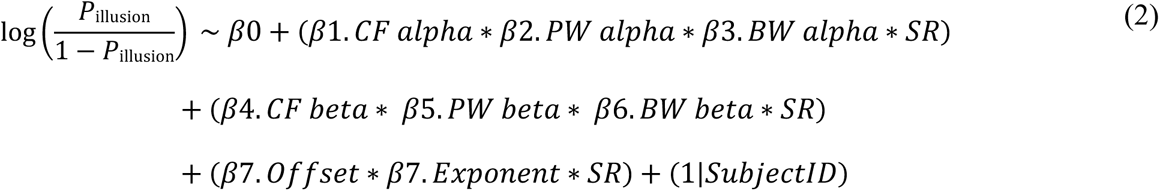

Where, P_illusion_ is the probability of perceiving the illusion, β_0_ is the intercept, β*n* are regression weights. The model constituted of continuous predictors (Offset, Exponent, CF (center frequency), PW (Peak parameter), and BW (bandwidth)) and categorical predictor SR (Sensor regions - factor of 5 levels: frontal, central, parietal, occipital, temporal) as interaction terms. Subject_ID was chosen as the random effect. Post-hoc analysis using Wald chi-square test were performed to get contrasts between the interaction terms using R package *car:Anova (type ‘III’)* (Fox J, Weisberg S, 2019).

### 2.7. Data and Code Availability

Raw EEG data used in the current study were analysed from a previously described study by Kumar *et al.,* 2020. For access to the raw data, kindly refer to the authors of the paper. However, processed data and relevant codes used for subsequent analysis in this study are available in the GitHub repository (https://github.com/VinseaSingh/Predicting-Response-to-McGurk-Illusion).

## 3. Results

### 3.1. Variability in illusory percept observed within and between participants^1^

Participants were subjected to four kinds of AV stimuli: incongruent McGurk (visual */ka/* paired with auditory */pa/* to induce the illusory percept */ta/*) and congruent (video and audio synched) */pa/*, */ta/*, */ka/*, presented at random. Participants were instructed to report whether they perceived */pa/*, */ta/*, */ka/*, or *"something else."*, unaware of the McGurk illusion. For each participant, total percentage response of the illusory percept /ta/ was calculated and reported as their McGurk susceptibility. We observed a broad spectrum of the total percentage responses for the eighteen participants with a median response at 60.83%. Nine of the eighteen participants showed <60.83% susceptibility towards the McGurk effect. And, the remaining nine participants showed >60.83% susceptibility towards the McGurk illusion (**Figure 1B** **(i)**).

We also calculated the response tendency (inter-trial variability) which is the relative proportion of responses for respective congruent and incongruent stimuli (Bechtold & Bastian, 2016; **Figure 1B** **(ii)**). Participants less prone to the illusion (<median McGurk susceptibility) on average reported an illusory /ta/ percept in 30.63% (SD = 19.83%) of trials whereas a non-illusory */pa/* percept was reported in 63.72% (SD = 22.35%) of trials, for the incongruent McGurk stimulus condition. Contrastingly, for participants more prone to the illusion (>median McGurk susceptibility) illusory /ta/ was perceived on average for 86.13% (SD = 10.89%) of the McGurk trials. And, non-illusory /pa/ was perceived for 11.06% (SD = 10.45%) of total trials. Congruent AV stimuli (*/pa/, /ta/, and /ka/*), in the case of less prone participants, were correctly identified in 96.02% (SD = 3.05%) of trials. Whereas for more prone, the congruent AV stimuli were reported in 97.05% (SD = 2.15%) of the trials. The difference between the percentage of */ta/* and */pa/* percept during the McGurk stimulus was significant for both the less prone (t_16_ = -3.32, p < 0.0043, Cohen’s d = -1.56) and more prone (t_16_ = 14.92, p < 0.001, Cohen’s d = 7.03) group of perceivers.

### 3.2. Differences in parametrized power was observed among participants before and after the illusory versus non-illusory responses to McGurk trials

The periodic and aperiodic component was extracted from trials that were sorted based on the perceptual categories (McGurk: /ta/ illusory and /pa/ non-illusory) for all the participants averaged across the sensors. This was done to generate a ‘global’ periodic and aperiodic parameter value representing the mean signal across the scalp across all participants. In case of periodic oscillations, illusory and non-illusory trails were statistically compared between subjects using cluster-based permutation independent-samples t-tests (Maris and Oostenveld, 2007). An observed t-value was permuted for 1000 iterations to generate the permutation distribution. Subsequently, the values of the observed cluster-level statistics were compared with the 5^th^ and 95^th^ quantile values of the respective permutation distribution. To counter for the multiple comparisons, we estimated the Tmax value at each iteration (Nichols & Holmes 2002). Spectral power in the theta (4-7 Hz), alpha (8-12 Hz), beta (15-30 Hz), and gamma (31-45 Hz) frequency bands were observed. We observed a lower alpha (t_5_ = -0.17) and lower beta band (t_5_ = -6.95) power for the illusory compared to non-illusory prestimulus trials (**Figure 2A**). Periodic oscillations of illusory trials post the McGurk stimulus showed a lower beta (t_5_ = -11.23) band activity as compared to non-illusory trials (**Supplementary Figure 2A**).

**Figure 2:**
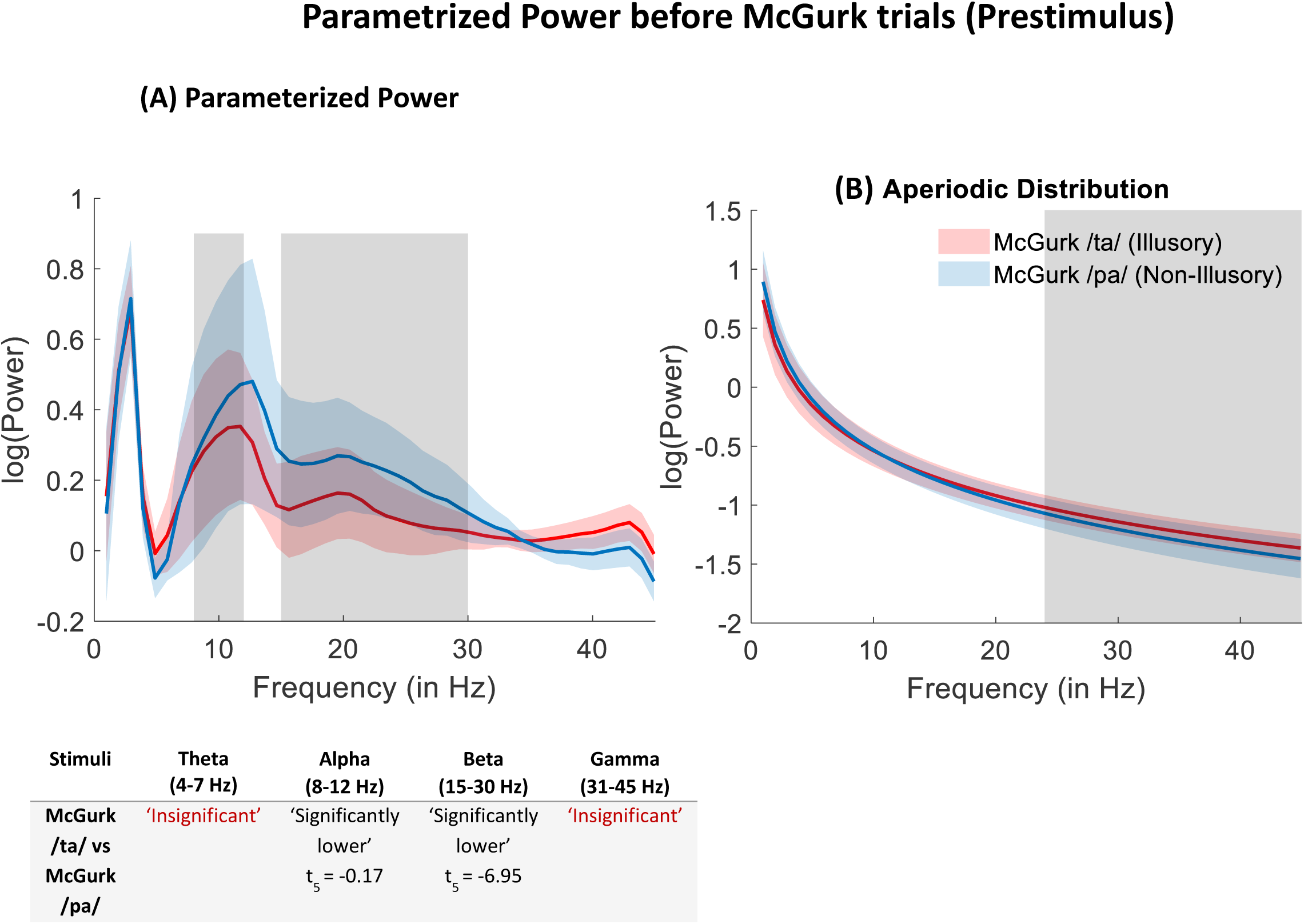
Parametrized power distributions before McGurk trials indicating inter-individual variability. (A) Prestimulus periodic power distributions averaged across all sensors for the illusory versus non-illusory McGurk trials. The table below indicates the frequency bands where the power was significantly different between the illusory (red) and non-illusory (blue) trials. (B) Prestimulus aperiodic distributions averaged across all sensors. The shaded region indicates the significant difference between illusory and non-illusory response condition (p = 0.042, two-tailed *t*-test) which is across 24-45 Hz frequency band.

The prestimulus aperiodic component sorted based on perceptual category were statistically compared using two-tailed t-test. We observed that aperiodic component before McGurk /ta/ response was higher than aperiodic component before McGurk /pa/ response in the frequency range between 24-45 Hz (t(44) = 2.09, p = 0.042, Cohen’s d = 0.62) (**Figure 2B**). And, for the aperiodic component post-illusory response was higher than in the frequency range between 7-45 Hz (t(78) = 1.99, p = 0.049, Cohen’s d = 0.47) as compared to post non-illusory response (**Supplementary Figure 2B**).

### 3.3. Periodic and aperiodic parameters showed significantly different sensor distribution before illusory and non-illusory McGurk trials

We estimated sensor-level distribution of prestimulus power parameters for before illusory and non-illusory McGurk response trials. For the periodic parameters, we looked in alpha and beta frequency ranges because we observed a significant difference at the global level. The statistical comparison across sensors were done using paired sample permutation t-test with observed t-value permuted for 1000 iterations and Tmax value was calculated. The prestimulus alpha periodic parameters before illusory percept, showcased a higher CF, PW, and BW in the frontal, central, parietal, and occipital sensors, and a lower activity in the temporal sensors (**Figure 3A**). Contrastingly, however, in the prestimulus beta periodic parameters, we observed a lower CF, PW, and BW in the frontal, central, parietal, and occipital sensors, and a higher activity in the temporal sensors before illusory percept (**Figure 3B**).

**Figure 3:**
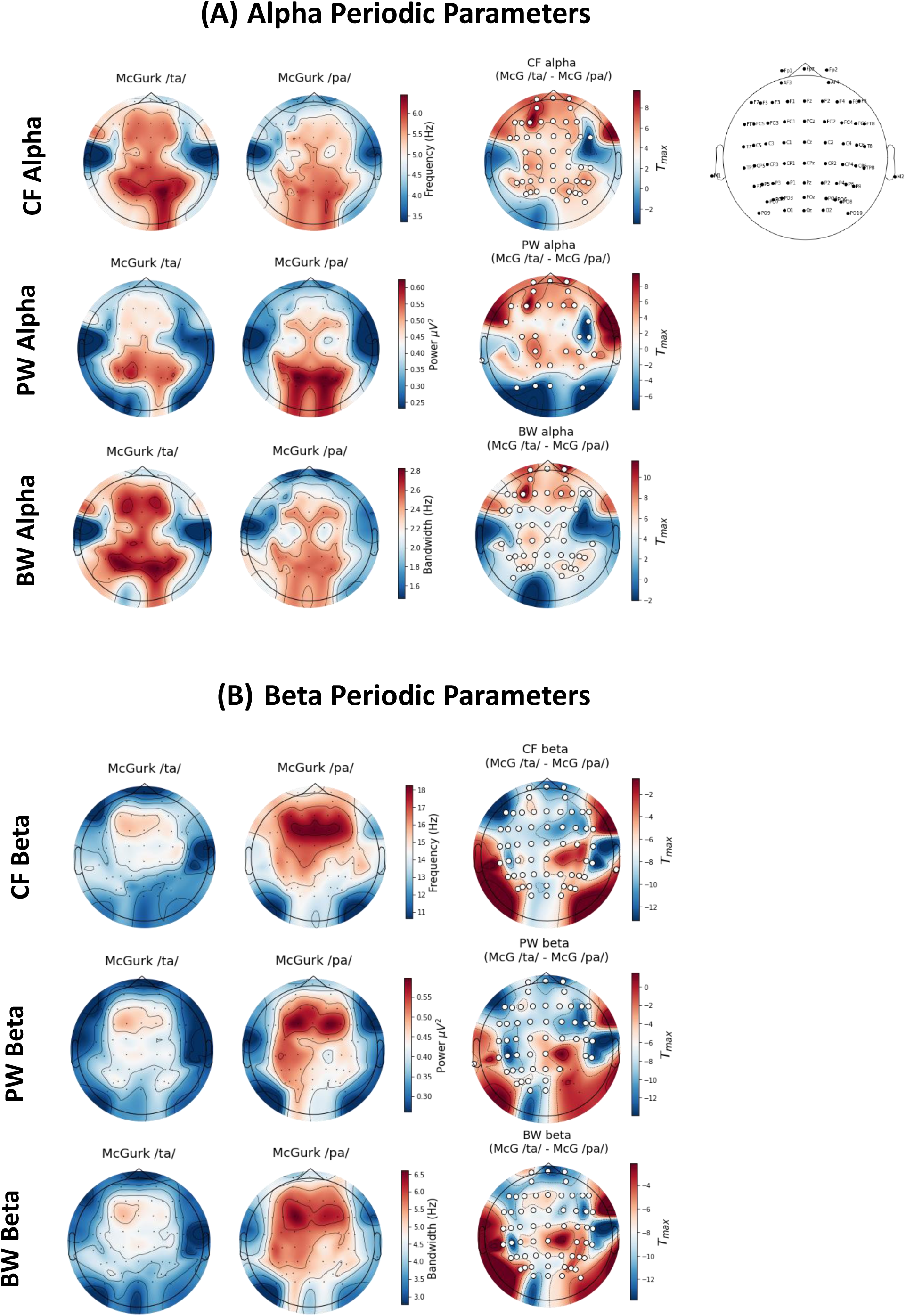
Topographies of prestimulus periodic power parameters (CF, PW, BW) for illusory (McGurk /ta/) versus non-illusory (McGurk /pa/) trials. (A) Topoplots of periodic parameters in the alpha (8-12 Hz) frequency range for the illusory, non-illusory response conditions and *t*-value maps (significant difference between the two conditions). White dots refer to the significant clusters of sensors with alpha at p < 0.05 (two-tailed). (B) Topoplots of periodic parameters in the beta (15-30 Hz) frequency range for the illusory, non-illusory response conditions and *t*-value maps (significant difference between the two conditions). White dots refer to the significant clusters of sensors with alpha at p < 0.05 (two-tailed).

In the aperiodic parameter offset, a higher activity before illusory percept was observed in the frontal, central, parietal sensors of the left hemisphere and parieto-occipital sensors of the right hemisphere, and lower activity in the temporal sensors, and right frontal and central sensors. Similarly for the aperiodic exponent, we observed a higher activity in the left frontal, central, and parietal sensors. And, a lower activity in the right frontal, right central sensors, temporal, and occipital sensors (**Figure 4**). Overall, these observations suggest a sensor cluster distribution of power parameters before illusory and non-illusory response to McGurk trials. And, to understand the effect of these parameters on the overall prediction of the response to the upcoming trial, we went ahead and grouped the electrodes into five different clusters covering bilateral frontal, central, parietal, occipital, and temporal sensors. See **Table 1** for more details.

**Figure 4:**
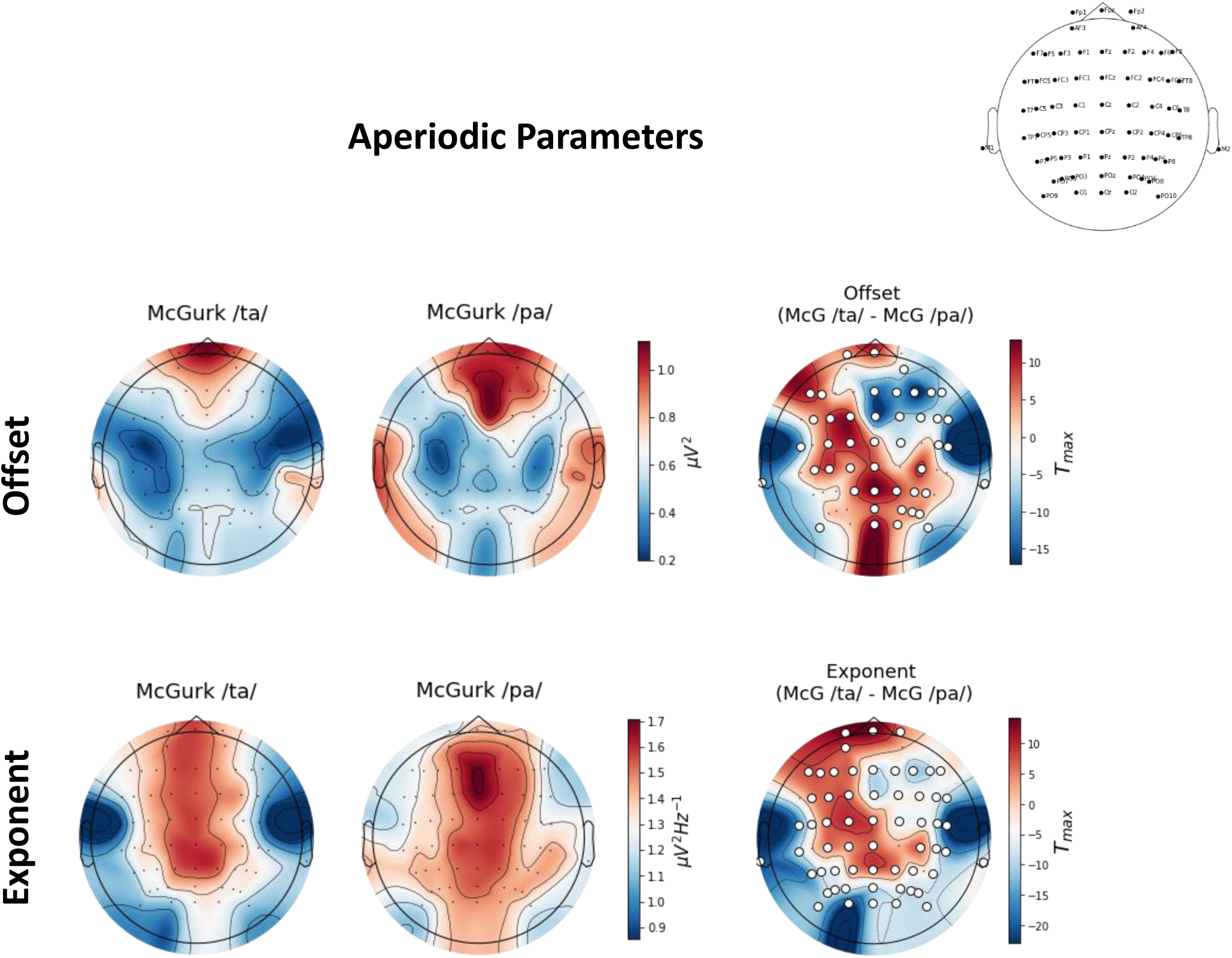
Topographies of prestimulus aperiodic power parameters (offset and exponent) for illusory (McGurk /ta/) versus non-illusory (McGurk /pa/) trials. Topoplots of aperiodic parameters offset (upper panel) and exponent (lower panel) for the illusory, non-illusory response conditions and *t*-value maps (significant difference between the two conditions). White dots refer to the significant clusters of sensors with alpha at p < 0.05 (two-tailed).

**Table 1:**
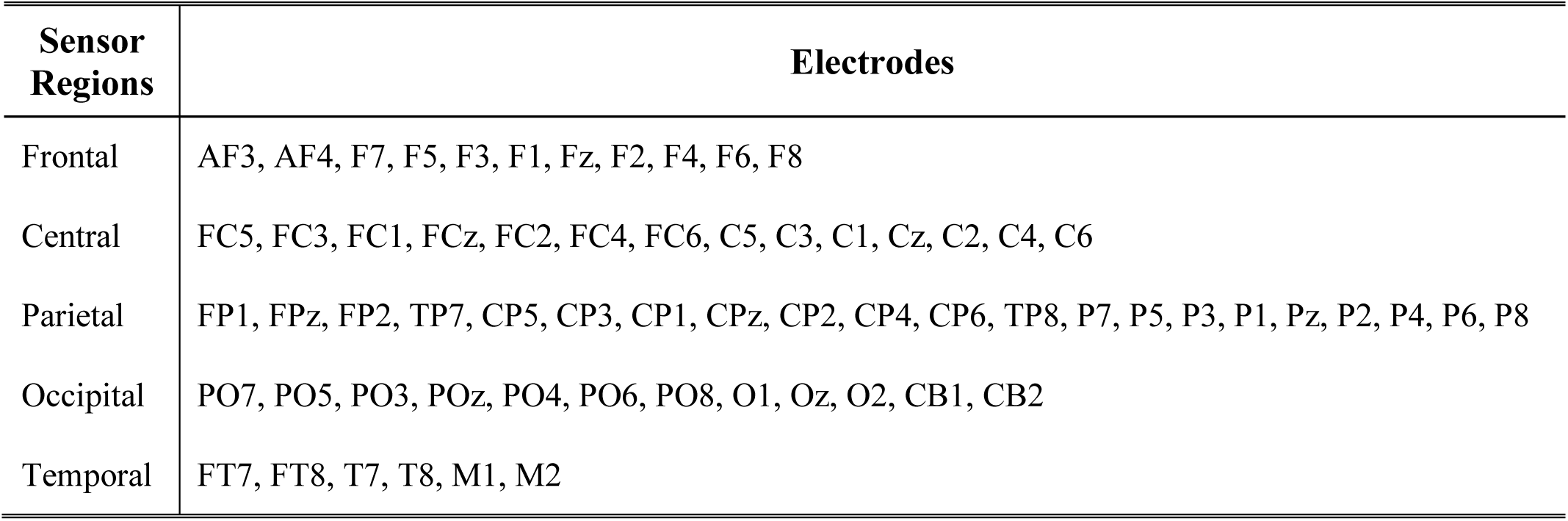
Table with the list of sensors categorized into respective sensor regions. We classified all the 64 sensors into five sensor regions covering bilateral: Frontal, Central, Parietal, Occipital, and Temporal.

| Sensor Regions | Electrodes |
| --- | --- |
| Frontal | AF3, AF4, F7, F5, F3, F1, Fz, F2, F4, F6, F8 |
| Central | FC5, FC3, FC1, FCz, FC2, FC4, FC6, C5, C3, C1, Cz, C2, C4, C6 |
| Parietal | FP1, FPz, FP2, TP7, CP5, CP3, CP1, CPz, CP2, CP4, CP6, TP8, P7, P5, P3, P1, Pz, P2, P4, P6, P8 |
| Occipital | PO7, PO5, PO3, POz, PO4, PO6, PO8, O1, Oz, O2, CB1, CB2 |
| Temporal | FT7, FT8, T7, T8, M1, M2 |

### 3.4. Periodic and aperiodic prestimulus parameters significantly differed before illusory and non-illusory perceptual trials at different sensor cluster

We estimated the differences between illusory and non-illusory trials at different sensor regions (**Table 1**) for the periodic (CF, BW, PW) and aperiodic parameters (Offset, Exponent). At the inter-individual level of variability, we observed significant differences for periodic oscillations in the alpha and beta frequency ranges, so we focused on the periodic parameters at these same frequency bands. The parameters were compared using Wilcoxon signed rank test, and p-values were adjusted using Holm-Bonferroni correction. The assumption for normality was assessed using Kolmogorov-Smirnov test. The test concluded that the data was non-normally distributed, justifying our choice of statistics. We also checked for the homogeneity of variances using Levene’s test where we obtained a p < 0.05 for all the periodic and aperiodic parametric distributions corroborating further on applying a non-parametric test.

We observed a significantly lower exponent before illusory perception across all sensor clusters (Central: Z = -3.789, p < 0.001, Cohen’s d = 0.061; Frontal: Z = -3.789, p < 0.001, Cohen’s d = 0.062; Occipital: Z = -7.033, p < 0.0001, Cohen’s d = 0.108; Parietal: Z = -2.452, p = 0.014, Cohen’s d = 0.037; Temporal: Z = -2.876, p = 0.004, Cohen’s d = 0.046) as compared to non-illusory perception for the McGurk trials, suggesting that ‘flatter’ prestimulus exponent might contribute globally to the illusory percept. Similarly, the offset was also observed to be significantly lower before illusory trials in all the sensor clusters except for the temporal region (Central: Z = -2.724, p = 0.006, Cohen’s d = 0.046; Frontal: Z = -2.573, p = 0.01, Cohen’s d = 0.042; Occipital: Z = -5.254, p < 0.0001, Cohen’s d = 0.083; Parietal: Z = -3.135, p = 0.002, Cohen’s d = 0.053) (**Figure 5A**).

**Figure 5:**
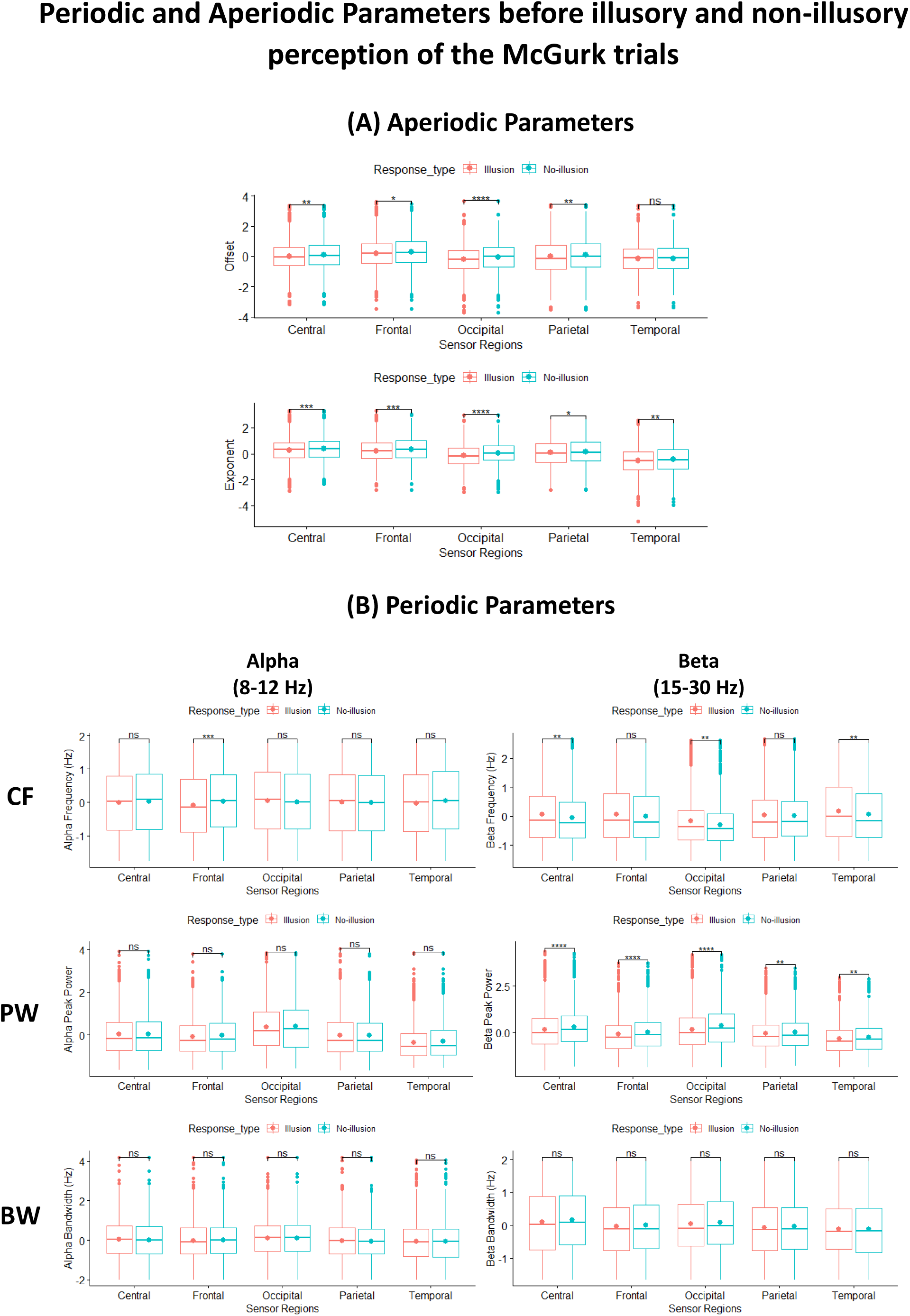
Boxplots comparing parameterized power values of illusory (red outline) versus non-illusory (blue outline) McGurk trials at different sensor regions using Wilcoxon signed rank test. (A) Aperiodic parameter (upper panel: Offset; lower panel: Exponent). (B) Periodic parameters (CF – peak frequency, PW – peak power, BW – peak bandwidth) in the alpha and beta frequency ranges. p-values were adjusted using Holm-Bonferroni correction, and asterisks above the boxplots describe the p-value: ‘*’ p ≤ 0.05, ‘**’ p ≤ 0.01 ‘***’ p ≤ 0.001, ‘****’ p ≤ 0.0001, and ‘n.s.’ non-significant.

For the prestimulus periodic parameters, a more intermittent differences were observed between the illusory and non-illusory trials across alpha and beta frequency bands (**Figure 5B**). Alpha center frequency was significantly different only at the frontal (Z = -3.746, p < 0.0001, Cohen’s d = 0.062) sensor clusters. The prestimulus beta center frequency was different between the two perceptual responses at the central (Z = -2.675, p = 0.0075, Cohen’s d = 0.046), occipital (Z = -2.675, p = 0.0075, Cohen’s d = 0.046), and temporal (Z = -2.689, p = 0.0071, Cohen’s d = 0.048) clusters. The aperiodic adjusted alpha peak parameter (PW), alpha bandwidth (BW), and beta bandwidth was observed to be insignificantly different between two response conditions across all sensor clusters. However, beta peak parameter was significantly lower for illusory conditions across all sensor clusters (Central: Z = -4.788, p < 0.0001, Cohen’s d = 0.075; Frontal: Z = -4.829, p < 0.0001, Cohen’s d = 0.076; Occipital: Z = -6.163, p < 0.0001, Cohen’s d = 0.096; Parietal: Z = -2.973, p = 0.003, Cohen’s d = 0.044; Temporal: Z = -3.270, p = 0.001, Cohen’s d = 0.052).

### 3.5. Periodic and aperiodic parameters of the prestimulus power can predict the response to the upcoming McGurk stimulus

We fitted a logistic mixed effect model (estimated using ML and BOBYQA optimizer) to predict the response to the upcoming McGurk stimulus with periodic parameters: center frequency, aperiodic adjusted power, and bandwidth in the alpha and beta frequency bands. And, aperiodic parameters: (offset and exponent) estimated for each sensor cluster (frontal, central, parietal, temporal, occipital) as the continuous predictors. The model included subject ID as random effect (formula: ∼1|Subject ID). The model’s total explanatory variance was moderate (conditional R^2^ = 0.21) and the variance of the fixed effects alone (marginal R^2^) was 0.095. The model’s intercept was at 0.79 (95% CI = [0.45, 1.12], p < 0.001). The intraclass correlation coefficient (ICC) of the model was 0.12, indicating that 12% of the variance of trail-wise response to the McGurk illusion depends on the subjects.

Moreover, within the model, we found significant effects of the certain predictors from different sensor regions. The model reported that the effect of prestimulus aperiodic exponent from frontal (β = - 0.20, 95% CI = [-0.37, -0.04], p = 0.015), central (β = -0.17, 95% CI = [-0.33, -0.02], p = 0.030), temporal (β = -0.27, 95% CI = [-0.40, -0.15], p < 0.001), and occipital (β = -0.37, 95% CI = [-0.53,-0.20], p < 0.001) sensor regions, and the effect of aperiodic offset from parietal (β = -0.19, 95% CI = [-0.34, -0.04], p = 0.016), temporal (β = 0.24, 95% CI = [0.12, 0.37], p < 0.001), and occipital (β = 0.18, 95% CI = [0.01, 0.35], p = 0.036) sensor regions could significantly predict the response to the upcoming McGurk stimulus. The model also reported that the periodic parameters like the alpha center frequency from frontal (β = -0.09, 95% CI = [-0.16, -0.03], p = 0.005), central (β = -0.07, 95% CI = [-0.14, -0.01], p = 0.038), and temporal (β = -0.09, 95% CI = [-0.15, -0.02], p = 0.011) regions; beta center frequency from the central (β = 0.10, 95% CI = [0.03, 0.17], p = 0.005), temporal (β = 0.10, 95% CI = [0.03, 0.17], p = 0.005), and occipital (β = 0.09, 95% CI = [0.02, 0.16], p = 0.013) regions; alpha peak parameter from parietal (β = 0.12, 95% CI = [0.04, 0.19], p = 0.003), and occipital (β = 0.09, 95% CI = [0.01, 0.18], p = 0.031) regions; aperiodic adjusted beta power from central (β = - 0.12, 95% CI = [-0.20, -0.04], p = 0.003), parietal (β = -0.12, 95% CI = [-0.19, -0.05], p = 0.001), and occipital (β = -0.11, 95% CI = [-0.19, -0.03], p = 0.005) regions could also significantly predict the response to the upcoming McGurk stimulus (**Table 2**).

**Table 2:** Logistic mixed-effect model of parameterized power parameters for predicting the McGurk illusory response across different sensor regions. The predictors are: CF alpha – peak alpha frequency, PW alpha – peak alpha power, BW alpha – peak alpha bandwidth; CF beta – peak beta frequency, PW beta – peak beta power, BW beta – peak beta bandwidth; Aperiodic parameter – Offset and Exponent. Predictors that significantly predicted the response are in bold.

| Predictors | Response |  |  | Bayes Factor |
| --- | --- | --- | --- | --- |
|  | Log-Odds | CI | p |  |
| (Intercept) | 0.79 | 0.45 – 1.12 | <0.001 | - |
| **<u>CF alpha</u>** |  |  |  |  |
| Frontal | -0.09 | -0.16 – -0.03 | 0.005 | 0.29 |
| Central | -0.07 | -0.14 – -0.00 | 0.038 | 3.10 |
| Parietal | 0.02 | -0.05 – 0.08 | 0.591 | 3.61 |
| Temporal | -0.09 | -0.15 – -0.02 | 0.011 | 1.34 |
| Occipital | 0.05 | -0.02 – 0.11 | 0.171 | 1.16 |
| **<u>PW alpha</u>** |  |  |  |  |
| Frontal | 0.01 | -0.07 – 0.09 | 0.782 | 4.42 |
| Central | 0.06 | -0.02 – 0.14 | 0.163 | 3.68 |
| Parietal | 0.12 | 0.04 – 0.19 | 0.003 | 1.26 |
| Temporal | 0.02 | -0.06 – 0.10 | 0.568 | 4.38 |
| Occipital | 0.09 | 0.01 – 0.18 | 0.031 | 0.51 |
| **<u>BW alpha</u>** |  |  |  |  |
| Frontal | -0.03 | -0.10 – 0.04 | 0.368 | 2.53 |
| Central | 0.06 | -0.01 – 0.13 | 0.113 | 3.81 |
| Parietal | 0.03 | -0.04 – 0.10 | 0.417 | 4.54 |
| Temporal | 0.03 | -0.03 – 0.10 | 0.325 | 3.84 |
| Occipital | -0.02 | -0.09 – 0.04 | 0.484 | 3.01 |
| **<u>CF beta</u>** |  |  |  |  |
| Frontal | 0.04 | -0.03 – 0.11 | 0.287 | 2.76 |
| Central | 0.10 | 0.03 – 0.17 | 0.005 | 0.08 |
| Parietal | -0.00 | -0.07 – 0.07 | 0.948 | 4.54 |
| Temporal | 0.10 | 0.03 – 0.17 | 0.005 | 0.19 |
| Occipital | 0.09 | 0.02 – 0.16 | 0.013 | 0.12 |
| **<u>PW beta</u>** |  |  |  |  |
| Frontal | -0.07 | -0.15 – 0.00 | 0.064 | 1.85 |
| Central | -0.12 | -0.20 – -0.04 | 0.003 | 0.77 |
| Parietal | -0.12 | -0.19 – -0.05 | 0.001 | 0.48 |
| Temporal | -0.04 | -0.11 – 0.03 | 0.315 | 4.34 |
| **Occipital** | **-0.11** | **-0.19 – -0.03** | **0.005** | **0.62** |
| ***BW beta*** |  |  |  |  |
| Frontal | -0.01 | -0.08 – 0.06 | 0.827 | 4.18 |
| Central | -0.04 | -0.11 – 0.03 | 0.304 | 1.99 |
| Parietal | -0.00 | -0.07 – 0.07 | 0.917 | 4.49 |
| Temporal | 0.00 | -0.07 – 0.07 | 0.938 | 4.22 |
| Occipital | 0.00 | -0.07 – 0.07 | 0.982 | 4.46 |
| ***Offset*** |  |  |  |  |
| Frontal | 0.13 | -0.05 – 0.30 | 0.148 | 4.50 |
| Central | 0.08 | -0.09 – 0.24 | 0.365 | 2.59 |
| **Parietal** | **-0.19** | **-0.34 – -0.04** | **0.016** | **0.08** |
| **Temporal** | **0.24** | **0.12 – 0.37** | **<0.001** | **0.16** |
| **Occipital** | **0.18** | **0.01 – 0.35** | **0.036** | 3.88 |
| ***Exponent*** |  |  |  |  |
| **Frontal** | **-0.20** | **-0.37 – -0.04** | **0.015** | 2.70 |
| **Central** | **-0.17** | **-0.33 – -0.02** | **0.030** | 4.18 |
| Parietal | 0.14 | -0.00 – 0.28 | 0.051 | **0.45** |
| **Temporal** | **-0.27** | **-0.40 – -0.15** | **<0.001** | **0.02** |
| **Occipital** | **-0.37** | **-0.53 – -0.20** | **<0.001** | **0.13** |
| **Random Effects** |  |  |  |  |
| $\sigma^2$ | 3.29 | | | |
| $\tau_{00}$ Subject_ID | 0.47 | | | |
| ICC | 0.12 |  |  |  |
| N Subject_ID | 18 |  |  |  |
| Observations | 4482 |  |  |  |
| Marginal R<sup>2</sup> / Conditional R<sup>2</sup> | 0.095 / 0.207 |  |  |  |
| AIC | 5711.673 |  |  |  |

A Bayes factor analysis established strong evidence of effects of central beta center frequency (BF = 0.08), parietal offset (BF = 0.08), and temporal exponent (BF = 0.02). The Bayes factor showed moderate evidence of effects of frontal alpha center frequency (BF = 0.29), temporal beta center frequency (BF = 0.19), temporal offset (BF = 0.16), occipital beta center frequency (BF = 0.12), and occipital exponent (BF = 0.13). And, finally the Bayes factor of central beta peak parameter (BF = 0.77), parietal beta peak parameter (BF = 0.48), parietal exponent (BF = 0.45), occipital alpha peak parameter (BF = 0.51), and occipital periodic beta peak parameter (BF = 0.62) showed weak evidence of effects on the overall model, meaning that these predictors had low impact on the prediction of response to McGurk trials. Interestingly, even though parietal exponent could not significantly predict the response (p = 0.051) it established anecdotal evidence of effects on the model fit (BF = 0.45). And finally, the Bayes factor, showed little to almost no effects of parietal periodic alpha power (BF = 1.26), temporal alpha center frequency (BF = 1.34), frontal exponent (BF = 2.70), central alpha center frequency (BF = 3.10), central exponent (BF = 4.18), and occipital offset (BF = 3.88) on the overall prediction model, even though these factors significantly predicted the response. See **Table 2** for more details.

### 3.6. A significant two-way and three-way interactions was observed among prestimulus periodic parameters in predicting response to the upcoming McGurk stimulus

We were interested in examining the effect of one independent predictor on another in predicting the response to the upcoming McGurk stimulus. We established interaction effects within periodic parameters of the alpha and beta frequency ranges and also among aperiodic parameters. To do so, we applied a logistic mixed effect model (estimated using ML and BOBYQA optimizer) on the interaction parameters (See Methods section for more details on the model). The model’s total explanatory power was moderate (conditional R^2^ = 0.25) and marginal R^2^ was of 0.02. The model’s intercept was at 0.92 (95% CI = [0.43, 1.41], p < 0.001). The intraclass correlation coefficient (ICC) of the model was 0.24, indicating that 24% of the variance of trail-wise response to the McGurk illusion depends on the subjects.

Post-hoc analysis revealed a combination of significant interaction effects for periodic parameters. A significant two-way interactions was observed between CF alpha x BW alpha (W(1) = 7.7392, p = 0.0054), and between PW beta x BW beta (W(1) = 4.1546, p = 0.0415). No significant two-way interactions were observed between CF alpha x PW alpha (p = 0.926), PW alpha x BW alpha (p = 0.748), CF beta x PW beta (p = 0.853), and CF beta x BW beta (p = 0.477). We also observed a significant interaction between CF alpha x Sensor Regions (W(4) = 10.5111, p = 0.0326). Adding all the periodic interactions together revealed a significant three-way interaction in the alpha band (CF alpha x BW alpha x PW alpha: W(1) = 6.8262, p = 0.0089) but not in the beta band (CF beta x PW beta x BW beta: p = 0.978).

For the aperiodic interaction parameters, however, we only observed two-way interactions between sensor regions and offset (W(4) = 16.4625, p = 0.0024), and also between sensor regions and exponent (W(4) = 18.4666, p = 0.001). No significant two-way interaction between Offset x Exponent (p = 0.469) and three-way interaction of Offset x Exponent x Sensor Regions (p = 0.546) was observed, indicating that the two aperiodic parameters independently predicted the response to the upcoming McGurk stimulus across different sensor clusters (**Table 3**).

**Table 3:** Fixed effects ANOVA (Type III Wald chi-square tests) results for logistic mixed-effect interaction model of parameterized power parameters. The predictors are: CF alpha – peak alpha frequency, PW alpha – peak alpha power, BW alpha – peak alpha bandwidth; CF beta – peak beta frequency, PW beta – peak beta power, BW beta – peak beta bandwidth; Aperiodic parameter – Offset and Exponent. Predictor interactions that significantly predicted the response are in bold.

| Predictors | Response |  |  |
| --- | --- | --- | --- |
|  | Chisq | DF | Pr (>Chisq) |
| **(Intercept)** | **13.7827** | **1** | **0.000205** *** |
| CF alpha | 3.003 | 1 | 0.083112 |
| PW alpha | 2.093 | 1 | 0.147975 |
| **BW alpha** | **3.8643** | **1** | **0.049323** * |
| Sensor Regions | 5.1596 | 4 | 0.271314 |
| **CF beta** | **6.9917** | **1** | **0.008189** ** |
| **PW beta** | **9.3843** | **1** | **0.002189** ** |
| BW beta | 1.2692 | 1 | 0.259911 |
| Offset | 0.8851 | 1 | 0.346798 |
| **Exponent** | **4.4705** | **1** | **0.034486** * |
| CF alpha x PW alpha | 0.0086 | 1 | 0.926291 |
| **<u>CF alpha x BW alpha</u>** | **7.7392** | **1** | **0.005404** ** |
| PW alpha x BW alpha | 0.1035 | 1 | 0.747678 |
| **CF alpha x Sensor regions** | **10.5111** | **4** | **0.032645** * |
| PW alpha x Sensor regions | 3.5639 | 4 | 0.468224 |
| BW alpha x Sensor regions | 3.5324 | 4 | 0.472963 |
| CF beta x PW beta | 0.0344 | 1 | 0.852834 |
| CF beta x BW beta | 0.5061 | 1 | 0.476817 |
| **<u>PW beta x BW beta</u>** | **4.1546** | **1** | **0.041521** * |
| CF beta x Sensor regions | 6.1927 | 4 | 0.185216 |
| PW beta x Sensor regions | 2.4133 | 4 | 0.66023 |
| BW beta x Sensor regions | 1.1293 | 4 | 0.889591 |
| Offset x Exponent | 0.523 | 1 | 0.469569 |
| **Offset x Sensor Regions** | **16.4625** | **4** | **0.002457** ** |
| **Exponent x Sensor Regions** | **18.4666** | **4** | **0.001** ** |
| **<u>CF alpha x PW alpha x BW alpha</u>** | **6.8262** | **1** | **0.008983** ** |
| CF alpha x PW alpha x Sensor Regions | 4.2194 | 4 | 0.377128 |
| CF alpha x BW alpha x Sensor Regions | 2.5545 | 4 | 0.634899 |
| PW alpha x BW alpha x Sensor Regions | 2.7414 | 4 | 0.601992 |
| CF beta x PW beta x BW beta | 0.0008 | 1 | 0.978147 |
| **CF beta x PW beta x Sensor Regions** | **10.8984** | **4** | **0.02773** * |
| CF beta x BW beta x Sensor Regions | 0.6933 | 4 | 0.952147 |
| PW beta x BW beta x Sensor Regions | 5.7238 | 4 | 0.220743 |
| Offset x Exponent x Sensor Regions | 3.0698 | 4 | 0.546207 |
| CF alpha x PW alpha x BW alpha x Sensor Regions | 3.7883 | 4 | 0.435411 |
| CF beta x PW beta x BW beta x Sensor Regions | 6.4656 | 4 | 0.166969 |

## 4. Discussion

The aim of this study was to investigate the relationship of single-trial prestimulus power and behavioral response to the upcoming McGurk stimulus. The prestimulus power spectral densities (PSDs) were parameterized to extract periodic and aperiodic components of the EEG signal. In case of McGurk illusory paradigms, the variability in activity is observed both at inter-individual and intertrial level (Nath & Beauchamp, 2011, 2012; Mallick *et al.,* 2015). So, we first established differences in the parameterized prestimulus power across participants (inter-individual variability) for illusory versus non-illusory conditions. We then determined sensor-wise distribution of the power parameters between the response conditions. And, finally using logistic mixed effect models, we estimated the neural signatures in the prestimulus duration that could possibly predict the response to the upcoming McGurk trial, independent of inter-individual variability. Several key findings emerged from this study. At the behavioral level, we observed a broad spectrum of responses to the illusion across all participants. We observed that people less prone to the illusion had higher percentage of non-illusory response as compared to people more prone to the illusory percept. This might be because less prone perceivers resolve the AV mismatch by focusing more on the auditory percept rather than the visual percept (here, lip movement articulating */ka/*), in case of multisensory stimuli (Morris *et al.,* 2017). Second, we observed a lower periodic alpha and lower periodic beta band oscillations for prestimulus activity before illusory versus non-illusory trails across all participants, along with significant broadband difference at 24-45 Hz frequency range in the aperiodic distribution for the two perceptual conditions. Third, sensor-wise distribution of prestimulus periodic and aperiodic parameters revealed positive and negative clusters between illusory and non-illusory McGurk trials. Fourth, we found prestimulus aperiodic parameter exponent to be significantly lower before illusory trials across all sensor regions. And finally, periodic and aperiodic component from different sensor regions successfully predicted the response to the upcoming McGurk stimulus, with periodic parameters eliciting significant two-way interactions within alpha and beta frequency ranges. And, a three-way interaction between alpha periodic parameters.

### 4.1. Decreased prestimulus alpha power and peak alpha frequency predicts illusory percept

Periodic or rhythmic neural oscillations are considered true oscillatory power, and they signify different cognitive or behavioral states (Engel *et al.,* 2001; Fries 2005; Cole & Voytek 2019). We observed a lower global alpha and lower global beta periodic power before illusory perception as compared to non-illusory trails across participants (**Figure 2A**). Keil and others have previously reported that only prestimulus beta power correlated with the perception of the McGurk illusion (Keil *et al*., 2012). However, we captured a significant alpha oscillatory difference between the response conditions. This might indicate that along with prestimulus beta, even alpha oscillations seem to modulate functional networks underlying illusion perception and that parametrizing the power spectrum brings out the effect of alpha activity in the perception of the McGurk illusion. Previous studies on simultaneity judgement tasks have reported a lower prestimulus alpha power when individual’s bias is towards visual input (Grabot *et al.,* 2017), leading to inaccurate simultaneity judgements of audio-visual stimuli (Bastiaansen *et al*., 2020). On similar lines, studies on McGurk illusion have reported that people fixating more on the lip movement tend to perceive the illusion (Gurler *et al*., 2015, Proverbio *et al*., 2016), so a lower prestimulus alpha observed in our study before illusory perception might predict participants’ bias towards the visual input. Another way to understand this bias is by looking at the literature on cortical excitability. Prestimulus alpha power have been inversely linked to cortical excitability especially in the visual cortex (Mathewson *et al.,* 2009; Romei *et al.,* 2008, 2010; Samaha *et al.,* 2017). Also, studies on stimulus detection tasks have reported that a lower prestimulus alpha oscillations indicate higher occipital excitability which leads to higher detection rates but lower accuracy (Benwell *et al.,* 2017, 2022; Lange *et al.,* 2014; Iemi *et al.,* 2017). Another set of literature reports prestimulus alpha power fluctuations to be linked to attention modulations, especially in the visual area (Haegens *et al.,* 2012; Jones *et al.,* 2010) and alpha oscillations not explicitly controlled (in our case prestimulus alpha power) could predict the response to multisensory stimuli. Our prediction model results align well with these studies. The logistic mixed-effect model showed that lower peak alpha power (PW) in the occipito-parietal regions predicted the response to illusory percept. This means that at regions with high-excitability (lower alpha power), the threshold of activation of the underlying neural population is lowered, thus improving the neural processing of sensory input leading to better perception, which in case of incongruent McGurk stimulus sometimes may lead to illusory percept (Lange *et al.,* 2014). Moreover, our model also showed that lower peak alpha frequency (CF) over frontal, central, and temporal electrodes predicted illusory response to the McGurk trial with stronger bayes factor evidence for frontal CF alpha (BF = 0.29). These results are contradictory to previous studies on sound induced flash illusion paradigms where they have reported that occipital individual peak alpha frequency predicts cross-modal integration of AV stimulus leading to illusory percept (Cecere *et al.,* 2015; Keil & Senkowski, 2017; Samaha & Postle, 2015; Venskus & Hughes, 2021). Overall, we propose that lower prestimulus alpha power and peak alpha frequency together modulate the shift in visual attention which might lead to McGurk illusory percept.

### 4.2. Decreased prestimulus beta power and peak beta frequency also predicts illusory percept

We observed a significantly lower beta power across all sensor regions; however, our model could only predict response to illusion perception from beta power over central, parietal and occipital electrodes. Our findings are consistent with previous research on audiovisual simultaneity judgement tasks where they showed that lower beta power over the scalp preceded visual-then-auditory sequences perceived as simultaneous trials (Yuan *et al.,* 2016). Prestimulus lower beta power have been associated with better sensory encoding (Griffiths *et al.,* 2019), which in case of incongruent AV stimulus input might lead to illusory perception (for review see, Keil & Senkowski, 2018). Recent rubber hand illusion studies, have shown reduced central alpha and beta band power before illusory percept (Evans & Blanke 2013; Rao & Kayser 2017). Interestingly, however, our results deviate from multisensory illusion studies where they have reported a higher prestimulus beta power before the illusory percept (Keil *et al.,* 2012; for review, see Keil & Senkowski, 2018; Kaiser *et al.,* 2019). These studies have looked at beta power without removing the *1/f* power (aperiodic) component, which might indicate that aperiodic component of the power spectrum majorly drives oscillations in the prestimulus duration and separating the periodic oscillations from aperiodic component brings out the true nature of oscillations (Monto *et al.,* 2008; Becker *et al.,* 2018; Donoghue *et al.,* 2020). Taking all these observations together, we propose that a lower true beta oscillation in the centro-parietal, and occipital regions drives McGurk illusory percept instead of the higher beta power. Moreover, we also observed that lower peak beta frequency (CF) in the central, temporal, and occipital regions predicted illusory responses. These peak beta frequencies might be modulating audio-visual attention shift along with peak alpha frequencies across different brain regions. However, looking at prestimulus cross-frequency coupling is beyond the scope of this study.

### 4.3. Decreased aperiodic parameter offset and exponent over the scalp predicts response to the illusory perception

The aperiodic (or arrhythmic) component of the power spectrum has been associated with modulations in cognitive states (Podvalny E. *et al.,* 2015; He B. *et al.,* 2010), aging (Voytek B. *et al.,* 2015; Donoghue *et al.,* 2020), and the excitation/ inhibition (E/I) balance of local neuronal populations (Manning *et al.,* 2009; Ray & Maunsell, 2011; Buzsaki *et al.,* 2012; Gao R. *et al.,* 2017; Waschke *et al.,* 2021). We found a significant decrease in the offset values and flatter slopes of 1/*f* background activity (exponent) before the illusory percept over the scalp. The lower prestimulus offset values from the parietal, temporal, and occipital electrodes could predict the illusory response to the upcoming McGurk stimulus, with the parieto-occipital sensors showing stronger evidence over the prediction model (BF = 0.08; BF=0.16). The offset parameter is referred to as signal’s baseline or "background noise," which is positively correlated to neural spiking (Manning *et al.,* 2009; Ray & Maunsell, 2011; Miller *et al*., 2012). Hence, our results on decreased offset value before illusory perception could reflect a decline in neuronal population spiking over the parieto-temporal sensors that lead to illusory percept.

Moreover, we also found that ‘flatter’ slopes over frontal, central, temporal, and occipital sensors could predict illusory response with stronger evidence seen in parietal (BF = 0.45), temporal (BF = 0.02), and occipital sensors (BF = 0.13). Exponent (or slope) parameter refers to rate of exponential power decay with increasing frequencies which is associated with underlying synaptic currents and reflects E/I balance. A flatter exponent reflects increased arrhythmic background neuronal firing which is thought to be driven by increased E/I ratio (Voytek *et al.,* 2015; Gao et al., 2017; Lendner *et al.,* 2020). However, the literature has inferred mixed understanding of this correlation in relation to cognitive processing. For instance, Wiltshire *et al.,* 2017 have reported that spectral slope becomes more ordered (or in this case steeper) with increasing demands on external attention. However, few studies have observed flatter slopes with higher state of consciousness (Miskovic *et al.,* 2019; Lendner *et al.,* 2020), and flatter slopes during rest to task state transitions (Podvalny *et al.,* 2015). Taking into account all of these different inferences, we propose that the effect of exponent is ’global,’ and whether the steeper or flatter slope defines cognitive processing is highly dependent on the nature of the stimulus input. In the case of multisensory McGurk stimulus input, a flatter slope in prestimulus duration from temporo-occipital sensors predicts response to the illusory percept.

### 4.4. Periodic parameters (CF, PW, BW) from the same frequency band greatly influence each other in predicting the illusory response but not aperiodic parameters (offset and exponent)

Our logistic mixed effect interaction model showed significant two-way interactions between peak alpha frequency, alpha bandwidth (alpha CF x alpha BW) and peak beta power, beta bandwidth (beta CF x beta BW), and a three-way interaction between peak alpha frequency, peak alpha power, and alpha bandwidth (alpha CF x alpha PW x alpha BW) for predicting the illusory response. Significant interactions means that the values of one independent predictor influences another’s overall performance in prediction (Fisher, 1992). These interactions for periodic parameters suggest that, while these different oscillatory features govern separate cognitive functions (Haegens *et al.,* 2014; Mierau *et al.,* 2017; Scally *et al.,* 2018), they influence each other’s effect on modulating prestimulus state, which leads to illusory or non-illusory percept in the case of McGurk stimulus. The nature of these interactions, however, has not been evaluated in this paper. Interestingly, we did not observe any significant interaction between aperiodic offset and exponent parameters in predicting the illusory response. Because both parameters define very distinct features of the power spectrum, they may not be influencing each other in the overall prediction of behaviour, and hence, we did not see any significant interaction between the two aperiodic parameters.

## 5. Conclusion

Perceptual variability during multisensory perception is determined by one’s ongoing brain state. In previous studies, these prestimulus brain states were characterized to capture oscillatory signatures that can predict one’s response to an upcoming McGurk stimulus, and it was found that higher beta power in lSTG in the prestimulus duration drives the illusory percept. In this study, we went a step further and isolated periodic and aperiodic power spectrum components from all sensors that predicted trial-wise response to an upcoming McGurk stimulus. Our logistic mixed effect model predicted the illusory response when we found periodic parameters (CF and PW) lower in fronto-central, parieto-occipital, and/or occipital sensors during the prestimulus duration. Lower parieto-temporal offset and ’global’ exponent values also predicted the illusory percept. In conclusion, our findings suggest that the prestimulus oscillatory state is majorly driven by aperiodic background activity, and that differences in these arrhythmic components cause inter-trial and inter-individual variability in McGurk illusion perception.

## Acknowledgements

This study was supported by NBRC Core funds, Ramalingaswami Fellowships (Department of Biotechnology, Government of India) to DR (BT/RLF/Re-entry/07/2014). DR was supported by SR/CSRI/21/2016 extramural grant from the Department of Science and Technology, Ministry of Science and Technology, Govt. India. Data used in the study was collected with funding from Department of Biotechnology, Ramalingaswami Fellowship (BT/RLF/Re-entry/31/2011) and Innovative Young Bio-technologist Award (IYBA), (BT/07/IYBA/2013) to AB. DR and AB acknowledge the generous support of the NBRC Flagship program BT/MEDIII/NBRC/Flagship/Program/2019: Comparative mapping of common mental disorders (CMD) over lifespan.

## Author contributions

**Vinsea AV Singh:** Conceptualization, Formal analysis, Methodology, Software, Visualization, Writing – Original draft preparation. **Vinodh G Kumar:** Investigation, Validation, Writing – Review & editing. **Arpan Banerjee:** Funding acquisition, Investigation, Project administration, Resources, Supervision, Validation, Writing – Review & editing. **Dipanjan Roy:** Conceptualization, Funding acquisition, Methodology, Project administration, Resources, Supervision, Validation, Writing – Review & editing.

## Conflict of interest

All the authors declare no conflict of interest.

## Supplementary Figure Captions

**Supplementary Figure 1:**
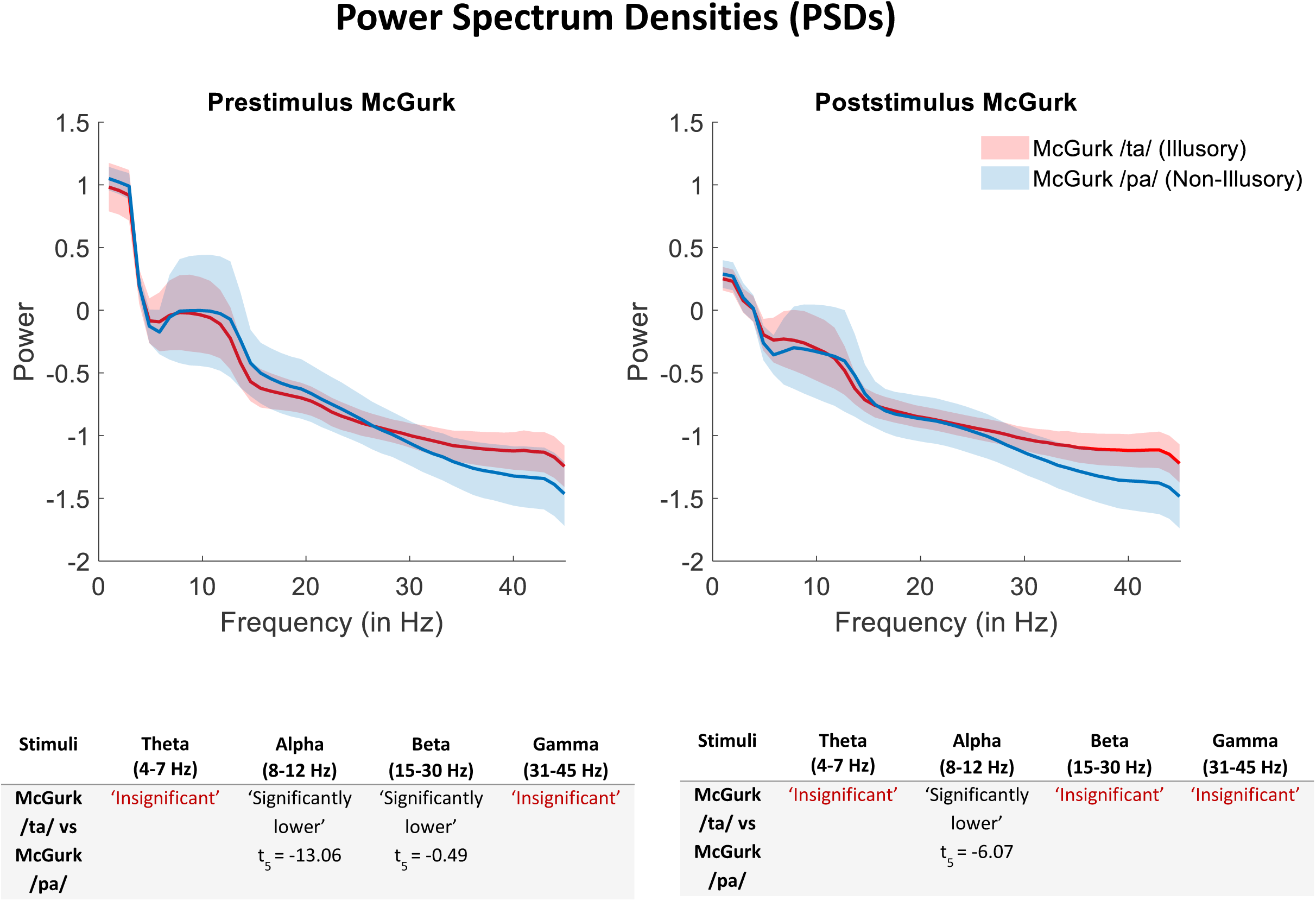
Power spectrum distributions before (left) and after (right) presenting McGurk trial. The spectra are averaged over all the sensors and across participants to capture the averaged activity difference between illusory and non-illusory McGurk perception. The table below each plot indicates the frequency bands where the power was significantly different between the illusory (red) and non-illusory (blue) trials. From the plots we can infer that in the prestimulus duration a significantly lower alpha and beta band power was observed for illusory trials. And, during the poststimulus duration we observed a lower beta band power after illusory perception.

**Supplementary Figure 2:**
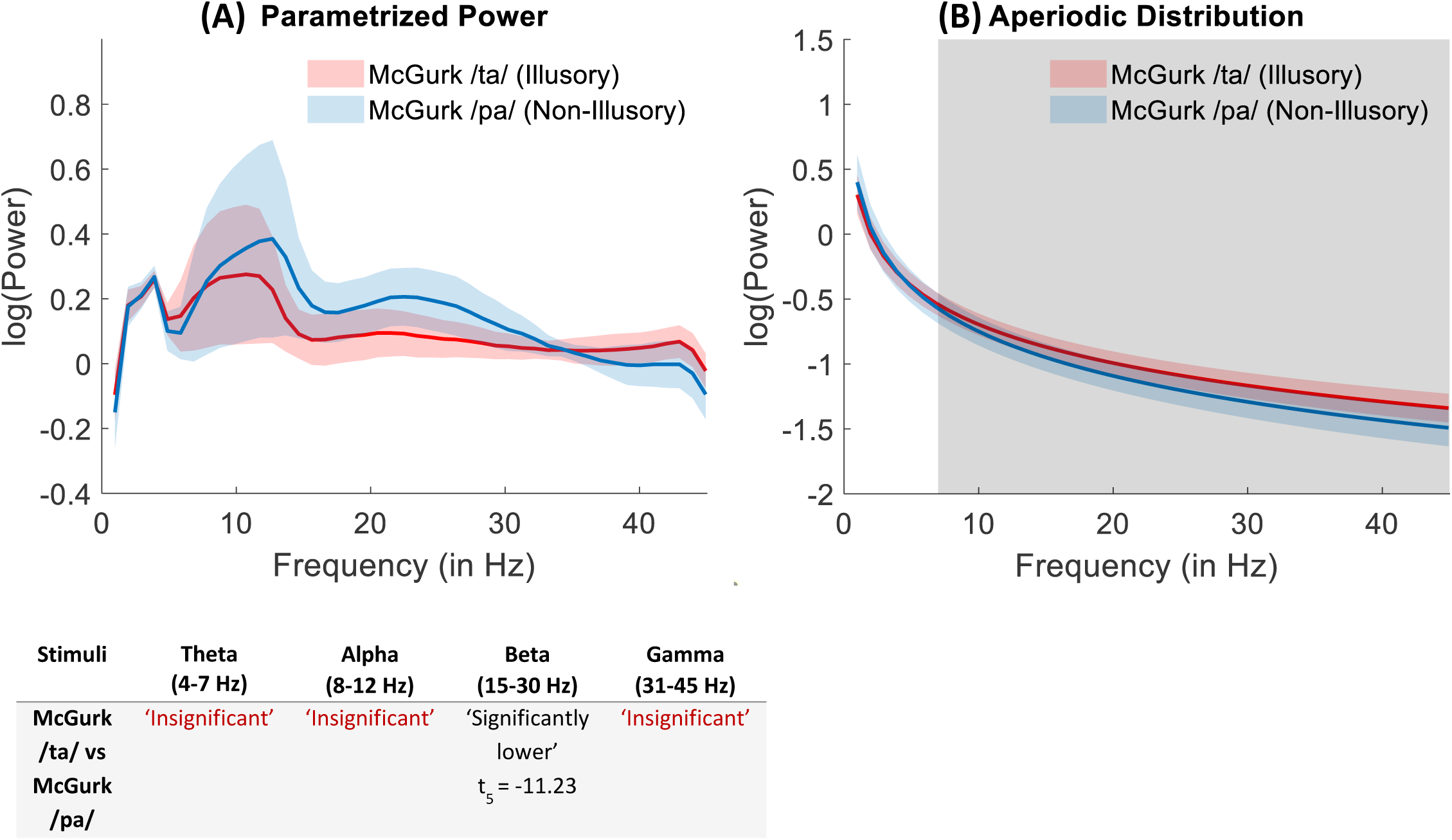
Parametrized power distributions after McGurk trials indicating inter-individual variability. (A) Poststimulus periodic power distributions averaged across all sensors for the illusory versus non-illusory McGurk trials. The table below indicates the frequency bands where the power was significantly different between the illusory (red) and non-illusory (blue) trials. (B) Poststimulus aperiodic distributions averaged across all sensors. The shaded region indicates the significant difference between illusory and non-illusory response condition (p = 0.049, two-tailed *t*-test) which is across 7-45 Hz frequency band.

## Footnotes

1 Data presented here were used in a different set of analyses by Kumar *et al.,* 2020.

## Notes

### Competing Interest Statement

The authors have declared no competing interest.

### Summary of Updates

In the revised version, the abstract, Introduction, methods, results, and discussion have undergone several modifications. All the figures and tables have been revised and modified substantially. The focus of this is now more on characterizing interindividual and intertrial variability of multisensory speech perception.

